# Identifying cross-lineage dependencies of cell-type specific regulators in gastruloids

**DOI:** 10.1101/2022.11.01.514697

**Authors:** Luca Braccioli, Teun van den Brand, Noemi Alonso Saiz, Charis Fountas, Patrick H. N. Celie, Justina Kazokaitė-Adomaitienė, Elzo de Wit

**Author notes:** Equal contribution (Luca Braccioli, Teun van den Brand).

## Abstract

Correct gene expression levels in space and time are crucial for normal development. Advances in genomics enable the inference of gene regulatory programs that are active during development. However, this approach cannot capture the complex multicellular interactions that occur during embryogenesis. Compared to model organisms such as fruit flies and zebrafish, the growth of mammalian embryos *in utero* further complicates the analysis of cell-cell communication during development. However, *in vitro* models of mammalian development such as gastruloids can overcome this limitation. Using time-resolved single-cell chromatin accessibility analysis, we have delineated the regulatory landscape during gastruloid development and thereby identified the critical drivers of developmental transitions. We observed that gastruloids develop from pluripotent cells driven by the transcription factor (TF) dimer OCT4-SOX2 and differentiate along two main branches. A mesoderm branch characterized by the TF MSGN1 and a spinal cord branch characterized by CDX1, 2, 4 (CDX). Consistent with our lineage reconstruction, ΔCDX gastruloids fail to form spinal cord. Conversely, *Msgn1* ablation inhibits the development of paraxial mesoderm, as expected. However, this also abolished spinal cord cells, which is surprising given that MSGN1 is not associated with differentiation along this branch. Therefore, formation of paraxial mesoderm is required for spinal cord development. To validate this, we generated chimeric gastruloids using ΔMSGN1 and wildtype cells, which formed both spinal cord and paraxial mesoderm. Strikingly, ΔMSGN1 cells specifically contributed to spinal cord, suggesting that cell-cell interactions between paraxial mesoderm and spinal cord are necessary for the formation of the latter. Our work has important implications for the study of cell-cell communication in development and how gene regulatory programs are functionally executed to form complex multicellular developmental structures.

## Introduction

Gastruloids are self-organizing, stem cell-based synthetic embryos that can reproducibly recapitulate key aspects of the early stages of post-occipital embryonic development. Gastruloids are generated by aggregating 300 mouse embryonic stem cells (mESCs) for 48 hours followed by a 24 hour pulse of the Wnt pathway agonist Chiron ^1^. Subsequently, the resulting cellular aggregates undergo germ layer specification and symmetry breaking, and can develop primitive organ structures such as spinal cord, somites, cardiac progenitors, blood and vascular tissue ^2–5^. Therefore, gastruloids are a valuable model to study development *in vitro* in a multi-tissue context. Individual cells within and across these tissues interact with each other physically and biochemically and the ensemble of these interacting cells can be viewed as a cell regulatory network (CRN) that orchestrates the emergence of tissue-level properties and determines the relationships between different tissues ^6,7^. Transcription within the individual cells of the CRN is driven by transcription factors (TFs), which act as central nodes inside gene regulatory networks (GRNs) that control key cellular processes such as cell differentiation ^8–12^. Linking GRNs, which are almost by definition cell intrinsic to cell-cell cooperation within a CRN is crucial for understanding developmental dynamics. Defining and perturbing the active TFs during gastruloid development may allow us to determine how alterations in the GRNs are propagated to the CRN level, and thereby advance our understanding of self-organization.

TFs are DNA binding proteins, but inside the context of the nucleus, nucleosomes can form a barrier for TF binding to gene regulatory elements. Therefore, the physical accessibility of these regions in chromatin acts as a regulatory mechanism ^13^. Differences in accessible chromatin regions can modulate the availability of regulatory elements for TF binding in a tissue-specific manner ^14,15^. Chromatin accessibility is established by pioneer TFs and chromatin remodeling complexes ^16^, paving the way for secondary TFs to bind these sequences. By measuring chromatin accessibility genome-wide by assay for transposase-accessible chromatin using sequencing (ATACseq) we can identify potential regulatory regions in a tissue. By analyzing the DNA binding motifs enriched in these regions it is possible to identify TFs associated with specific cell identities during gastruloid development ^14,17–19^.

In order to capture the gene regulatory complexity within developing gastruloids, we resorted to time course single cell combinatorial indexing ATACseq (sciATACseq) ^20^ and delineated the main regulatory subtypes in mouse gastruloids. We found that these subtypes strongly match their embryonic counterpart. By studying TF DNA binding motif enrichment along the regulatory trajectories of the cell types we identified the TFs CDX and MSGN1 as drivers of the early cell fate decisions generating the two main developmental branches (spinal cord and paraxial mesoderm respectively). In particular, we determined that *Cdx* genes promote neuromesodermal progenitor (NMP) differentiation and spinal cord development, but are dispensable for mesoderm formation. Moreover, we found that ablation of MSGN1 causes impairment of paraxial mesoderm differentiation as well as cross-lineage inhibition of spinal cord formation. To discriminate between the intrinsic and extrinsic roles of MSGN1 we combined genetic deletion with chimeric gastruloid formation and found that in chimeric gastruloids MSGN1 null cells contribute efficiently to spinal cord formation. Altogether our work represents a novel framework for delineating GRNs and determining how perturbations to the GRN are propagated to the CRN and affect gastruloid development.

## Results

### sciATACseq identifies regulatory cell fate transitions during gastruloid development

To define the gene regulatory landscape guiding gastruloid development, we probed a modified version of sciATACseq ^20^ (see Methods) on gastruloids derived from E14 mouse embryonic stem cells (mESC) and collected at 72 (i.e. 24h post chiron treatment), 80, 88, 96 and 120h (Figure 1A-B). We also included gastruloids that were cultured in 5% Matrigel between 96 and 120h, which develop into trunk-like structures (TLS) ^4^ (Figure 1A-B). We obtained high quality chromatin accessibility data for 12,575 nuclei with a median of 18,845 unique fragments per nucleus and an average fraction of reads in peaks (FriP) of 0.336 (Figure S1A-F). To determine whether clearly defined subtypes exist, we performed UMAP visualization of the single cell accessibility data (Figure 1C). Cluster analysis identified 8 cell subtypes originating from a progenitor population which differentiate along a temporal trajectory (Figure 1C). The proportion of progenitor cells diminishes sharply from 72h onward and essentially disappears at 96h (Figure 1D). We detected a cluster likely representing endoderm/gut which comprised very few cells. These cells were excluded from the rest of the analysis on the basis of them being very rare (Figure 1C).

**Figure 1:**
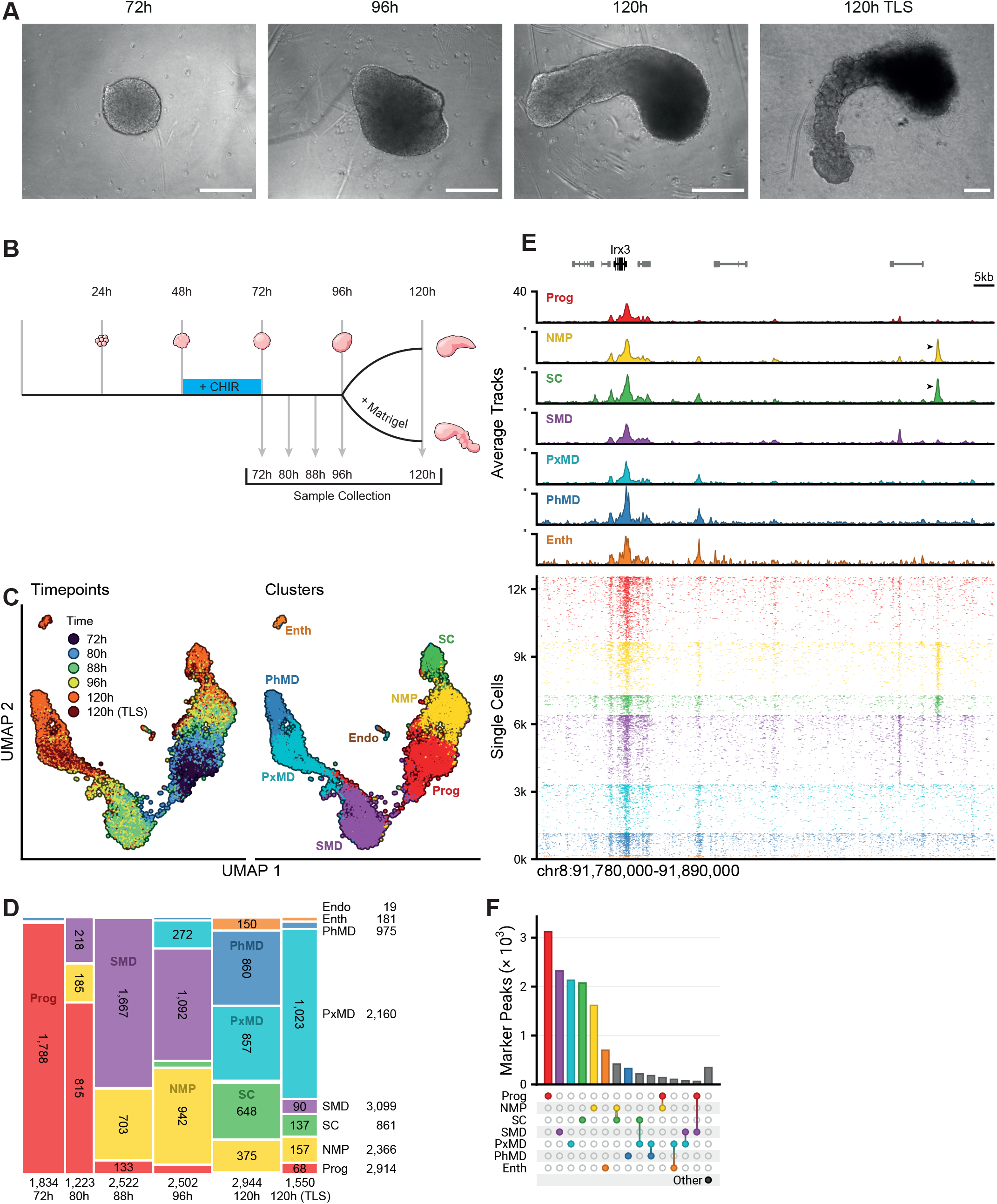
A chromatin accessibility map of gastruloid development. A) Bright-field microscopy images of typical gastruloids at 72, 96 and 120 hours into the culturing protocol. The 120 hour trunk like structure (TLS) was taken after culturing gastruloids in 5% matrigel from 96 hours onwards. Scale bar: 200 µm. B) Schematic overview of the culturing protocol and sample collection strategy for sciATAC-seq. C) Dimensionality reduction through UMAP for single cell accessibility data, coloured by collection time-points (left) or after Leiden clustering (right). Clusters are labelled by inferred cell types. Prog: progenitors; NMP: neuromesodermal progenitors; SC: spinal cord; Endo: endoderm; SMD: somitic mesoderm; PxMD: paraxial mesoderm; PhMD: pharyngeal mesoderm; Enth: endothelium. D) Example locus near the Irx3 gene, displaying pseudo-bulk coverage by cluster at the top and single cell coverage at the bottom. Arrows indicate a region specifically accessible in the NMP and SC clusters. E) Mosaic plot displaying areas proportional to the cell numbers for combinations of collection time-points and clusters. F) Upset plot giving the number of unique marker peaks per cluster and frequent combinations of clusters. The ‘Other’ category gives the sum of all marker peaks that were too infrequent to be included.

In Figure 1E we show chromatin accessibility for individual cells and as aggregate signal for the different clusters nearby the developmentally important *Irx3* gene ^21^. We observed regions that are ubiquitously accessible across all clusters, which often coincide with promoter regions (e.g. the promoter of the *Irx3* gene). On the other hand, we found distal regulatory regions that are accessible only in a subset of clusters (Figure 1E). We observe that the vast majority of promoter-proximal peaks are not differential during gastruloid development (Figure S1H). To determine cell type specific regulatory elements, we defined accessible chromatin regions specific to each subpopulation. To this end we calculated the differentially accessible (DA) peaks for each cluster and will be referred to as marker peaks (Figure 1F).

### Gastruloid regulatory landscapes strongly resemble *in vivo* counterparts

For gastruloids to be a *bona fide* model for developmental gene regulatory networks the regulatory landscapes in gastruloids should resemble *in vivo* regulatory landscapes. To determine this we compared our dataset with sciATACseq data generated from embryonic day E8.25 mice ^14^, which is right after gastrulation and at the beginning of organogenesis. Similarities between early embryos and gastruloids were calculated by integrating the nuclei from our dataset with nuclei from the embryo dataset and made quantitative predictions using support-vector machine (SVM) ^22^. We found a high degree of similarity between most gastruloid subpopulations and the embryo cells (Figure 2A). We used the SVM predictions to assign labels to gastruloid cells (Figure S2A). We identified gastruloid subtypes corresponding to NMPs, spinal cord cells, paraxial mesoderm, pharyngeal mesoderm, somitic mesoderm and endothelium. The majority of these cell types are found in post-occipital tissues, with the exception of pharyngeal mesoderm which is an anterior tissue ^23^. Among the populations detected, we identified two intermediate cell types. Somitic mesoderm represents an intermediate state which is strongly reduced at 120h, after giving origin to paraxial mesoderm at 96h and pharyngeal mesoderm at 120h (Figure 1E) ^24^. Similarly, the NMP pool, known to give rise to spinal cord cells *in vivo* ^25^, is reduced at 120h when spinal cord cells increase (Figure 1E). We failed to convincingly predict a corresponding embryo cell type for the progenitor cluster (Figure 2A), therefore we speculated that these cells are similar to the starting mESC population. Indeed, marker peaks for the progenitor cluster show specific enrichment in bulk ATACseq data from E14 mESCs (Figure 2B). To orthogonally confirm our labeling, we plotted pseudobulk chromatin accessibility of the embryo sciATACseq data Figure 2B). This shows that cell-type specific accessible regions in gastruloids are also accessible in the embryo counterparts. These patterns are further exemplified by the chromatin accessibility landscape near known cell type-specific genes (Figure 2C). The cell types that we identified are consistent with the cell types found in gastruloid scRNAseq data ^3^. Taken together, these analyses indicate that gastruloids contain several regulatory subpopulations which faithfully recapitulate important aspects of the post-occipital embryonic regulatory landscape.

**Figure 2:**
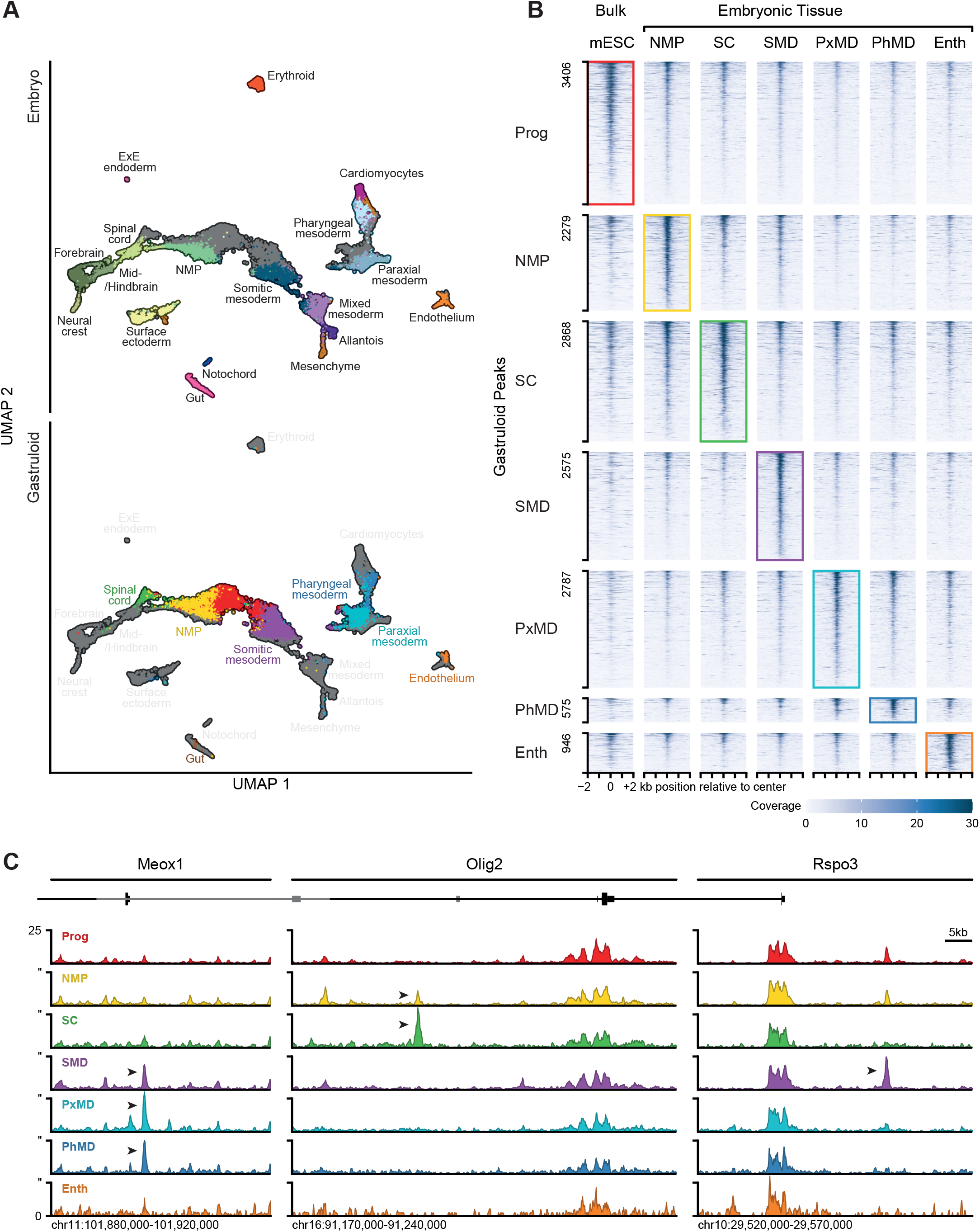
Gastruloid accessibility mirrors accessibility in several embryonic tissues. A) Dimensionality reduction through UMAP for the combined single cell accessibility data from embryos ^14^ and gastruloids. Top displays embryo cells by cluster and gastruloid cells in grey. Bottom displays gastruloid cells by cluster and embryo cells in gray, with well-overlapping tissues coloured in black. B) Tornado plot showing pseudo-bulk accessibility coverage of tissues in the embryo ^14^ and bulk coverage in WT mESCs at sites identified as markers for gastruloid populations. C) Pseudo-bulk coverage tracks of gastruloid data at loci nearby cell-type specific genes. Notable differentially accessible sites are highlighted with arrows.

### Chromatin state transitions highlight developmental trajectories in gastruloids

Our time-resolved sciATACseq dataset enables us to determine the regulatory cascade underlying gastruloid development. To this end we performed pseudotime analysis, which orders cells along the progression of an imputed differentiation trajectory ^26^. We identified three main pseudotime trajectories starting from the progenitor cells: I) NMP-spinal cord, II) somitic mesoderm-paraxial mesoderm and III) somitic mesoderm-pharyngeal mesoderm. The paraxial and pharyngeal mesoderm trajectories consist of a common root originating at the progenitor population and transitioning through the somitic mesoderm population (Figure 3A). These trajectories corroborate that NMPs give origin to spinal cord ^25^. However, in contrast with the established origin of somitic mesoderm from NMPs in mouse embryos, our pseudotime analysis suggests that in gastruloids, mesoderm cells can be directly derived from the progenitor cluster ^25^ (see Discussion for more details). It should be noted that we could not determine a trajectory for the endothelial cluster and therefore also no endothelial precursor cells. The developmental trajectories that we identified can be used as a starting point to define the regulatory cascade contributing to cell fate specification and morphogenesis.

**Figure 3:**
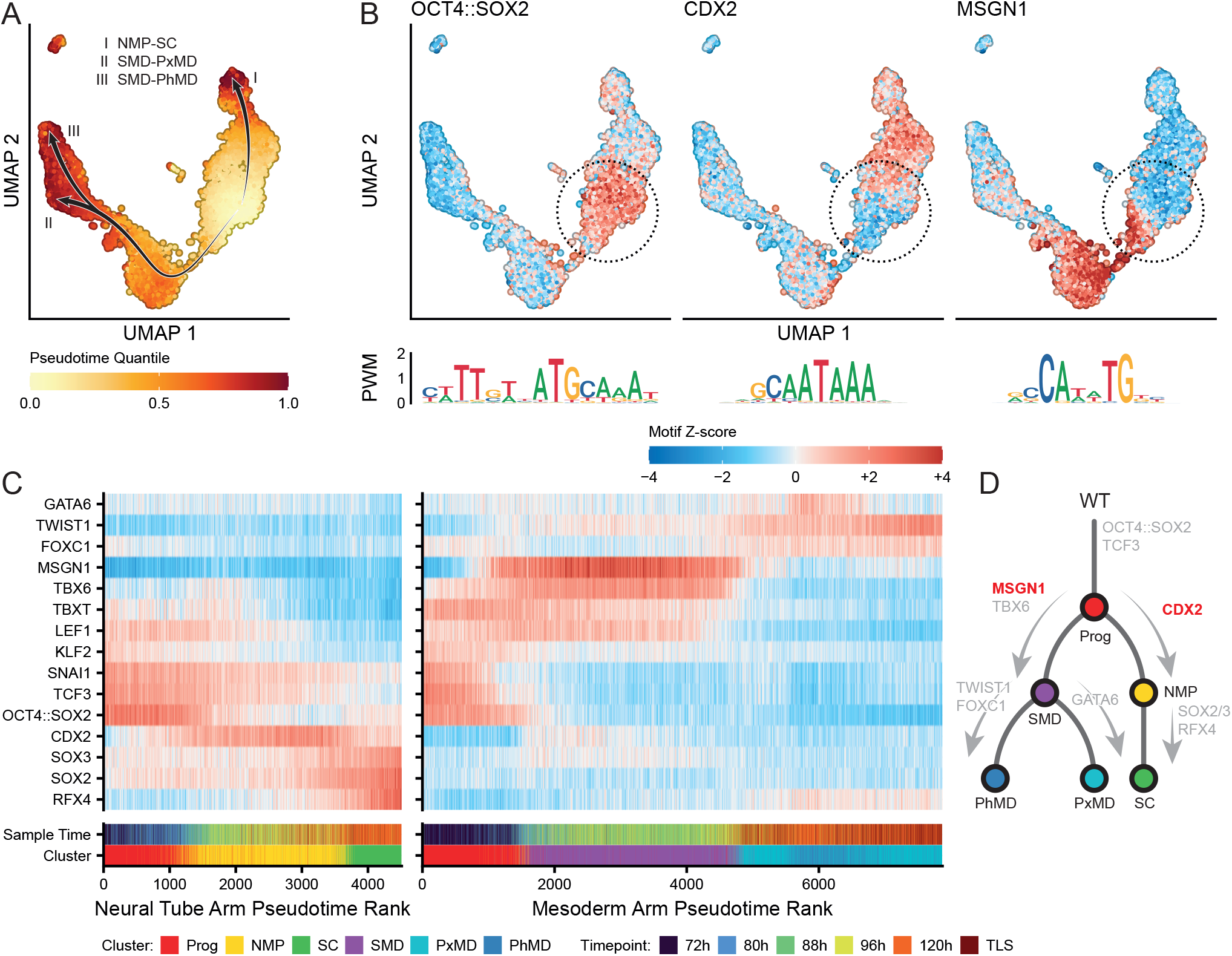
CDX2 and MSGN1 motifs are transiently enriched while progressing through gastruloid development. A) Pseudotime indicated in UMAP plot with proposed branches of differentiation indicated with arrows. B) UMAP plot displaying cell-wise enrichment of transcription factor binding site motifs for the OCT4::SOX2 compound motif, CDX2 and MSGN1, along with their position weight matrices displayed as logos. C) Heatmap displaying motif enrichment in cells ordered along the proposed (I) NMP-spinal cord branch of differentiation (left) and (II-III) somitic–paraxial/pharyngeal mesoderm branch (right). Every cell represents a binned average of 10 cells. Motif Z-scores are computed as GC-bias corrected measures describing gain or loss of accessibility at sites where the motifs were detected relative to the population’s average.

### TF motif accessibility along cell trajectories reveals potential drivers of cell fate decisions

The changes in the regulatory landscape during gastruloid development are largely driven by TFs. To identify key driver TFs binding to cell type specific regulatory elements, we performed DNA binding motif enrichment analysis on the DA peaks. We found that in the progenitor cluster regulatory elements were enriched for the pluripotency-associated motif OCT4::SOX2, bound by OCT4 and SOX2, and binding sites for the Wnt effector TF TCF3 ^27,28^ (Figure 3B-C). TCF3 binding to the OCT4::SOX2 motif in ESCs has been suggested to compete with SOX2 binding and counteract pluripotency, indicating that the cells in the progenitor cluster are undergoing differentiation ^29^. Accordingly, a subset of the progenitor cells shows enrichment of the binding motif of mesoderm marker TBXT (a.k.a. Brachyury) ^30^. Next, to identify the TFs that potentially drive subpopulation development we calculated TF motif enrichment along the pseudotime trajectories. We decided to focus on the two main branches we identified: the NMP-spinal cord and the somitic mesoderm-paraxial mesoderm trajectory (Figure S3A).

On the NMP-spinal cord trajectory, we observed a decrease in the enrichment of the OCT4::SOX2 and TCF3 motifs as the progenitor cells transition to the NMP state (Figure 3B-C). The binding motif of CDX2 (highly overlapping with CDX1 and CDX4 motifs) is highly enriched in the NMP population. Interestingly, we find the CDX2 motif also enriched in a subset of cells within the progenitor population (Figure 3B-C).

This indicates that these cells are already primed for differentiation towards the NMP-spinal cord trajectory. We found that the levels of CDX2 motif enrichment decreased again upon differentiation of NMPs into spinal cord cells (Figure3B-C), pattern indicating that CDX1, 2, 4 TFs may be specifically required in the NMP stage and no longer necessary once the cells differentiate further. Supporting this hypothesis, it has been shown that *Cdx* genes maintain the NMP state by suppressing the retinoic acid signaling pathway which is required for differentiation of NMPs into spinal cord cells ^31,32^. When we re-analyzed gastruloid scRNAseq for the expression of *Cdx1, -2* and *-4* in NMPs, we found that in the majority of NMPs *Cdx2* was expressed at a higher level than *Cdx1* and *-4* (Figure S3B) ^4^, consistent with the observation that CDX2 plays a prominent role in posterior axis elongation ^33,34^. During the transition from NMPs to spinal cord, we detected an increase of SOX2 and RFX4 motif enrichment (Figure 3B, C). In line with the motif analysis, RFX4 is expressed in spinal cord cells in gastruloids (Figure S3B). RFX4 has been shown to promote ventral spinal cord development in mice by inducing Sonic Hedgehog (Shh) signaling to ensure primary cilia formation and represents a marker of spinal cord differentiation in gastruloids ^35^. SOXB family members have a binding motif highly similar to that of SOX2 and have been implicated in neural differentiation and in spinal cord differentiation ^36,37^. In gastruloids spinal cord cells express SOX1, -2, -3 and -21 ^4^ (Figure S3B). SOX2 in particular is known to promote the exit from the NMP stage and the differentiation into spinal cord cells ^31,38,39^.

Next, we analyzed the TF motifs found in the somitic mesoderm-paraxial mesoderm trajectory. Similar to the NMP-spinal cord trajectory, we observed the OCT4::SOX2 and TCF3 motif enrichment decreasing abruptly when transitioning from progenitors to the somitic mesoderm and acquiring enrichment for the binding motif of MSGN1 (Mesogenin1, Figure 3B-C). Along this trajectory, we already detect an enrichment of the MSGN1 motif within a proportion of the progenitor cells, suggesting that these progenitors become primed for differentiation (Figure 3B-C). When the cells enter the paraxial mesoderm state, we see that MSGN1 and TBX6 motif enrichment is rapidly decreased, consistent with a role for MSGN1 and TBX6 that is restricted specifically to somitic mesoderm ^40,41^ (Figure 3C). Moreover, MSGN1 is known to promote the transition between mesoderm progenitor cells and presomitic mesoderm ^31^. TBX6 has been reported to induce MSGN1 expression which could explain why TBX6 enrichment precedes MSGN1 motif enrichment ^40^ (Figure 3C). We also found an enrichment for the motif of the epithelial-to-mesenchymal transition (EMT) TF TWIST1 in the paraxial mesoderm cells (Figure 3C). In the embryo, EMT is crucial to transition from the initial primitive streak epithelial cells towards paraxial mesoderm ^42^. When we determined TF motif enrichment dynamics across cellular states we did not observe sharp transitions (except for the endothelial cluster), but rather a continuous modulation of the level of accessibility of these motifs (Figure 3B, C). These observations suggest that cells do not suddenly acquire a differentiated phenotype but rather transition from a progenitor state to an intermediate state, where TFs associated with either progenitor or differentiated states bind chromatin simultaneously in the same cells. In conclusion, the analysis of TF binding motifs along the developmental trajectories allowed us to identify relevant lineage-specific TF activities and infer their dynamics during cell fate specification (Figure 3D). Moreover, we found that priming for differentiation occurs already at the (intermediate) progenitor states.

### *Cdx* genes drive posterior elongation by inducing NMP differentiation into spinal cord but not into paraxial mesoderm

Given the transient gain of open chromatin regions enriched for the CDX motif in the NMP-spinal cord branch we wanted to test the functional requirement of CDX1,2,4 in the formation of spinal cord cells during gastruloid development. To this end we attempted to generate gastruloids from ΔCDX1,2,4 (ΔCDX) HM1 mESCs ^31^. Compared to gastruloids formed from wild-type (WT) HM1 mESCs, the ΔCDX mESCs failed to form elongated structures during the 120h of the gastruloid differentiation protocol (Figure 4A-B). Although the observed structures are lacking some key features of gastruloids, we will refer to them as gastruloids for the sake of clarity and on the basis of that they went through the gastruloid protocol. The observed phenotype mimics the effect of *Cdx* deletions in mouse embryos, which leads to a failure in posterior axis elongation ^33,43^. Based on our pseudotime trajectory we would predict decreased formation of SOX2-positive spinal cord cells (Figure 3A-C). Indeed, ΔCDX gastruloids displayed reduced expression of the spinal cord marker SOX2 when compared to WT gastruloids (Figure 4C-D). Moreover, we did not observe differences in expression of the mesoderm marker FOXC1 ^44^ in ΔCDX gastruloids compared to WT gastruloids, suggesting that ΔCDX gastruloids are still capable of forming paraxial mesoderm (Figure 4C-D). Additionally, we observed that TBXT expression, partially co-localizing with CDX2 expressing cells at the posterior end of WT gastruloids, was diminished and diffused throughout ΔCDX gastruloids (Figure S4A-B). Taken together, differentiation in the ΔCDX structures is diverted from the spinal cord towards the mesoderm.

**Figure 4:**
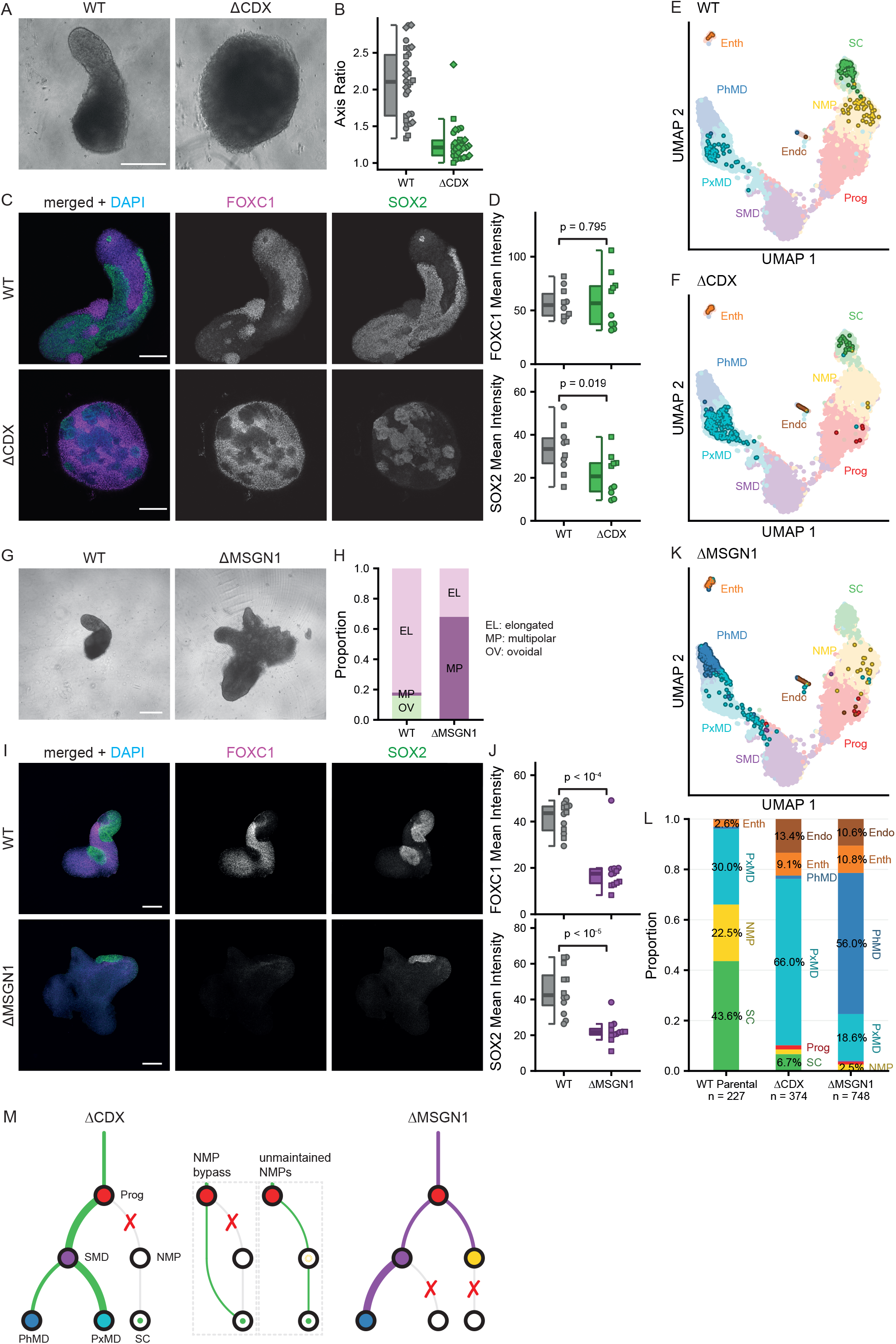
Effect of CDX and MSGN1 deletion on gastruloid cell type composition. A) Bright-field microscopy images of WT and ΔCDX gastruloids at 120 hours. Scale bar: 200 µm. B) Axis ratio quantification of WT and ΔCDX gastruloids (n = 3 biological replicates; rep. 1 = circles, WT: 10 gastruloids, ΔCDX: 8 gastruloids; rep. 2 = squares, WT: 10 gastruloids, ΔCDX: 10 gastruloids; rep. 3 = diamonds, WT: 10 gastruloids, ΔCDX: 12 gastruloids). C) Confocal immunofluorescence (IF) images of WT and ΔCDX gastruloids at 120 hours, stained for the FOXC1 mesoderm marker (magenta) and SOX2 spinal cord marker (green). DAPI in blue. Scale bar: 200 µm. D) Quantification of FOXC1 and SOX2 expression in WT and ΔCDX gastruloids at 120 hours (n = 2 biological replicates; rep. 1 = circles, WT: 5 gastruloids, ΔCDX: 5 gastruloids; rep. 2 = squares, WT: 5 gastruloids, ΔCDX: 5 gastruloids). E) UMAP plots highlighting cells from WT gastruloids at 120 hours relative to cells obtained from the time series experiment displayed in muted colours. F) Same as E but for ΔCDX gastruloids. G) Bright-field microscopy images of WT and ΔMSGN1 gastruloids at 120 hours. Scale bar: 200 µm. H) Quantification of morphology of WT and ΔMSGN1 gastruloids (n = 3 biological replicates; rep. 1: WT: 10 gastruloids, ΔMSGN1: 10 gastruloids; rep. 2: WT: 20 gastruloids, ΔMSGN1: 20 gastruloids; rep. 3: WT: 20 gastruloids, ΔMSGN1: 20 gastruloids). I) Like C but for WT and ΔMSGN1 gastruloids at 120 hours. J) Quantification of FOXC1 and SOX2 expression in WT and ΔMSGN1 gastruloids at 120h (n = 2 biological replicates; rep. 1 = circles, WT: 7 gastruloids, ΔMSGN1: 6 gastruloids; rep. 2 = squares, WT: 5 gastruloids, ΔMSGN1: 6 gastruloids). K) Same as E and F but for ΔMSGN1 gastruloids. L) Quantification of population proportions found in WT, ΔCDX and ΔMSGN1 gastruloids at 120h. M) Schematic summary of the differentiation trajectories measured in WT, ΔCDX and ΔMSGN1 gastruloids.

Although the immunofluorescence analysis is useful for measuring the spatial distribution of marker genes it cannot determine the composition of regulatory subtypes in the ΔCDX gastruloids. Therefore, we performed sciATACseq at 120h in WT gastruloids and ΔCDX gastruloids. WT HM1 gastruloids displayed a cell-type composition similar to our E14 WT 120h gastruloids, indicating that our sciATACseq and gastruloid protocols are robust and reproducible (Figure 4E). ΔCDX gastruloids displayed a near absence of NMPs and a strong reduction of spinal cord, consistent with our immunofluorescence results (Figure 4F). Additionally, we observed a strong increase of paraxial mesoderm cells. This shows that the impairment in spinal cord specification is compensated by the acquisition of a paraxial mesodermal fate. Importantly, this confirms our prediction that paraxial mesoderm in gastruloids does not derive from NMPs.. These observations strengthen the finding that *Cdx* genes act in a trajectory-specific manner by promoting the formation of NMPs and, by extension, spinal cord cells. However, *Cdx* genes are not required for mesoderm differentiation. This suggests that, in gastruloids, NMPs are not a bipotent progenitor pool but they contribute solely to spinal cord development. In summary, *Cdx* genes are required for spinal cord formation but not for paraxial mesoderm differentiation.

### MSGN1+ cells are required for spinal cord formation

Because we observed an enrichment of the MSGN1 motif in the somitic mesoderm-paraxial mesoderm pseudotime trajectory, we investigated the role of MSGN1 during gastruloid development. We performed gastruloid differentiation using ΔMSGN1 mESCs ^31^ which yield non-elongating structures when compared to WT cells at 120h (Figure 4G-H). Moreover, we observed that ΔMSGN1 gastruloids presented multiple CDX2+/TBXT+ protrusions (Figure S4C, E). Based on our pseudotime analysis, we predicted MSGN1 to promote mesoderm formation (Figure 3A-C). The drastic reduction in FOXC1 expressing cells in ΔMSGN1 gastruloids when compared to WT gastruloids confirmed these predictions (Figure 4I-J). Unexpectedly, we also observed a reduction in SOX2+ spinal cord cells in ΔMSGN1 gastruloids (Figure 4K-L). sciATACseq analysis of ΔMSGN1 gastruloids showed a massive shift of paraxial mesoderm to pharyngeal mesoderm (barely detected in WT cells) (Figure 4E, K, L). This suggests that MSGN1 is promoting paraxial mesoderm identity while suppressing the emergence of pharyngeal mesoderm, consistent with *Msgn1*^-/-^ mutant mouse embryos exhibiting loss of paraxial mesoderm ^45,46^. To our surprise, the sciATACseq did not detect any cells with a regulatory landscape reflecting spinal cord cells in ΔMSGN1 gastruloids. Additionally, we observed a strong reduction in the percentage of NMPs. This contrasted with our prediction, since the MSGN1 motif is depleted in the cells that form the spinal cord trajectory (Figure 3). We conclude that within the developing gastruloids there are dependencies between the trajectories that are not readily apparent from our sciATACseq analysis. We hypothesize that defects in spinal cord formation in ΔMSGN1 gastruloids result from a defect in a cross-lineage mechanism involving paraxial mesoderm and spinal cord cells that guides neural fate acquisition (Figure 4H).

We reasoned that if the differentiation of NMPs to spinal cord cells is independent of MSGN1, but rather is dependent on the interaction between different cell types, differentiation from NMPs to spinal cord cells should occur when this interacting environment is extrinsically provided. Gastruloids are the ideal model system for this as it is possible to mix different mESC lines to create chimeric gastruloids ^47^. This approach provides a unique opportunity to dissect lineage extrinsic regulatory pathways. To test whether ΔMSGN1 could form spinal cord cells when provided with the correct cellular environment we generated chimeric gastruloids composed of an equal mix of WT and ΔMSGN1 mESCs. To be able to track WT and ΔMSGN1 cells, we labeled the cells with mCherry and EGFP, respectively. Since labeling the HM1 mCherry+ WT line failed to reproducibly form gastruloids (data not shown), we resorted to E14 WT mCherry+ cells for the following experiments. Note that HM1 and E14 are both derived from 129 mice (see method section for details).

In contrast to the gastruloids derived from ΔMSGN1 EGFP+ cells only, chimeric gastruloids consisting of WT mCherry+ and ΔMSGN1 EGFP+ cells were elongated at 120h (Figure 5A-B). In fact, elongation rates in chimeric gastruloids were similar to WT gastruloids (Figure 5B). This shows that in chimeras, WT mCherry+ cells can compensate for the lack of MSGN1 in half of the initial cell pool and restore gastruloid elongation. Strikingly, in the chimeric gastruloids, ΔMSGN1 EGFP+ cells were enriched at the posterior end of the chimeric gastruloids, while WT mCherry+ cells were distributed along the anterior (Figure 5A, C)..

**Figure 5:**
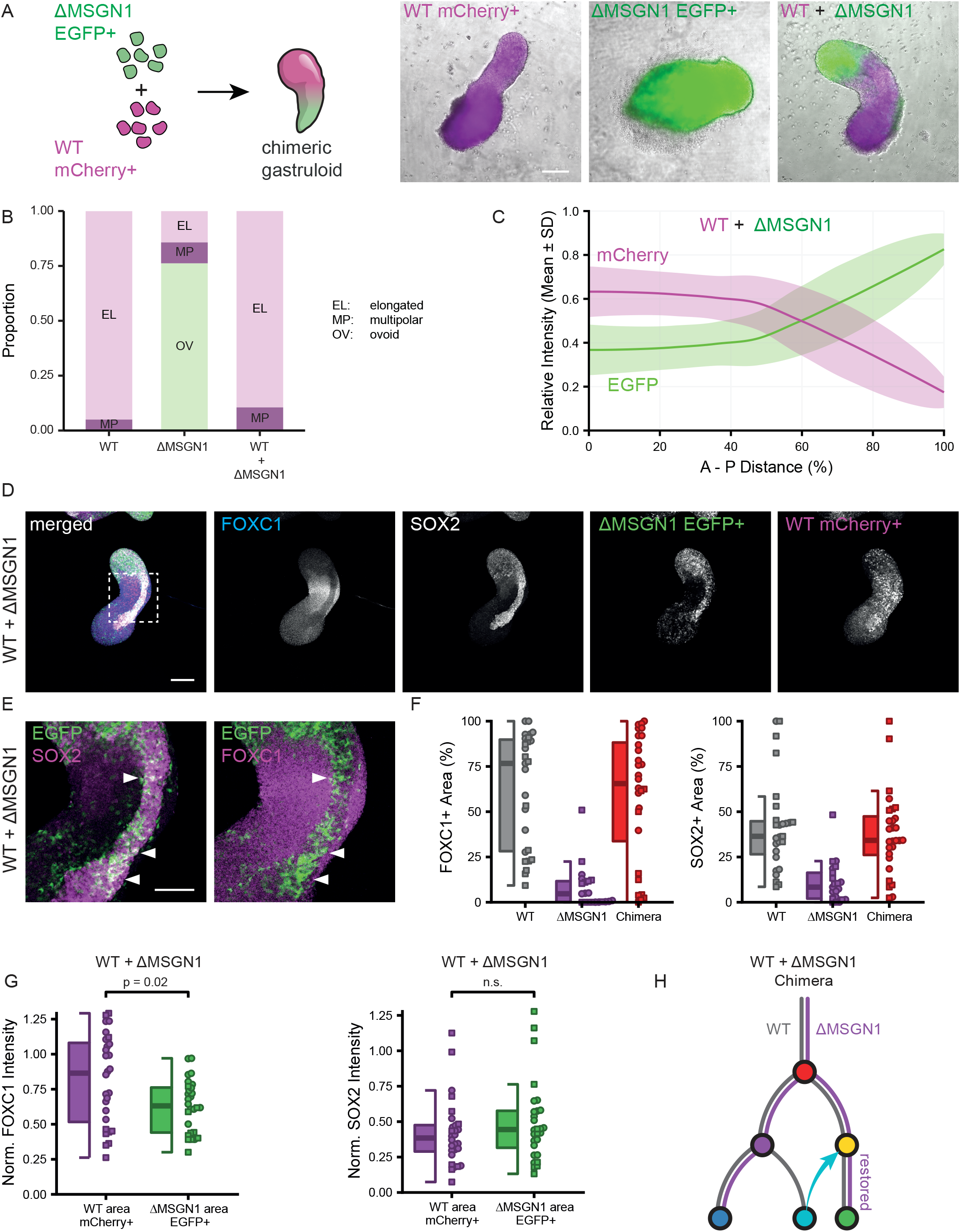
ΔMSGN1 cells can form spinal cord cells in presence of WT cells that competently form paraxial mesoderm. A) Schematic representation of chimeric gastruloids and wide-field fluorescence microscopy images of fluorophore expressing WT only, ΔMSGN1 only and chimeric WT + ΔMSGN1 gastruloids at 120 hours. Scale bar: 200 µm. B) Quantification of morphology of WT, ΔMSGN1 and chimeric gastruloids (n = 2 biological replicates; rep. 1: WT: 10 gastruloids, ΔMSGN1: 10 gastruloids, chimeras: 10 gastruloids; rep. 2: WT: 10 gastruloids, ΔMSGN1: 11 gastruloids. chimeras: 9 gastruloids). C) Quantification of fluorophore intensity in chimeric gastruloids along the AP axis. D) Confocal immunofluorescence image of WT + ΔMSGN1 chimeric gastruloids at 120 hours, staining for FOXC1 (blue), SOX2 (grey) and EGFP (green) and mCherry (magenta). Scale bar: 200 µm. E) Detail of (D) indicated by the dashed square, showing ΔMSGN1 cells contributing to spinal cord formation. SOX2 and FOXC1 in magenta as indicated, EGFP in green. Scale bar: 200 µm. F) Quantification of FOXC1 and SOX2 expression in WT, ΔMSGN1 and chimeric gastruloids. G) Normalized quantification of FOXC1 and SOX2 expression in mCherry+ (WT) and EGFP+ (ΔMSGN1) areas in chimeric gastruloids. The intensities over all optical sections were summarised per gastruloid by taking a weighted mean of intensity, where the area in each section acted as the weight. The mean intensity of each replicate was normalized on the mean intensity of the corresponding WT sample. H) Schematic summary of differentiation trajectories measured in chimeric gastruloids.

To determine the contribution of WT mCherry+ and ΔMSGN1 EGFP+ cells to mesoderm and spinal cord, we performed immunofluorescence analysis of lineage markers. We did not observe multiple CDX2+/TBXT+ protrusions in ΔMSGN1 EGFP+ gastruloids as opposed to our experiments with the untagged cells (Figure S4C, E; Figure S5A). We think this could have resulted from repeatedly subculturing the ΔMSGN1 mESCs in order to achieve EGFP tagging of the cells. Compared to WT mCherry+ gastruloids, ΔMSGN1 EGFP+ gastruloids showed a strong reduction in FOXC1+ mesoderm as well as SOX2+ spinal cord, consistent with our previous observations (Figure 5 D, F; Figure S5B). Importantly, chimeric gastruloids displayed SOX2+ spinal cord and FOXC1+ mesoderm regions comparable to WT mCherry+ gastruloids (Figure 5D, F; Figure S5B). In the chimeras, we observed both FOXC1+ and SOX2+ positive cells derived from the WT mCherry+ mESCs, meaning that they contributed to both spinal cord and mesoderm formation (Figure 5D, G). However, more importantly, we observed that in chimeras ΔMSGN1 EGFP+ cells expressed SOX2+ but displayed lower levels of FOXC1 (Figure 5D-E, G). This indicates that the presence of mesoderm formed by WT cells rescues the capacity of ΔMSGN1 cells to form spinal cord cells, and therefore provides strong evidence of a cell fate induction mechanism that promotes spinal cord identity (Figure 5H). Together with the fact that in chimeras ΔMSGN1 EGFP+ did not form FOXC1+ mesoderm efficiently, we have shown that MSGN1 is cell autonomously required for mesoderm formation, but is required in a cross-lineage manner for spinal cord differentiation. Taken together these data provide evidence that the paraxial mesoderm appears to be a critical guiding component in order for NMPs to progress to spinal cord.

## Discussion

Here, we systematically dissect the chromatin regulatory landscape during mouse gastruloid development, defining the main regulatory subtypes that arise in this synthetic embryo model. We find that besides the initial progenitor pool, the majority of the subpopulations have a high degree of resemblance to their counterparts in post-gastrulation embryos. This resemblance suggests that gastruloids faithfully recapitulate cell types and associated regulatory landscapes in posterior embryo development, complementing the findings of transcriptomic studies ^3,4^. For most of the identified subpopulations, we were able to define clear developmental trajectories stemming from the progenitor cluster. However, we did not find a trajectory explaining the emergence of the endothelial population, which was found almost exclusively found at 120h in our sciATACseq dataset. This late emergence suggests that their origin may be pinpointed by including additional time points between 96 and 120h. However, a recent transcriptomic study indicates that endothelial cells derive from mesoderm ^48^. When analyzing TF DNA binding motif enrichment along the main developmental trajectories, we observed that from each (intermediate) progenitor state to the next population, the cells enter a “bivalent” state, i.e. display enrichment for accessibility of the motifs of OCT4::SOX2 as well as of either MSGN1 or CDX2, suggesting that progenitors undergo regulatory priming before differentiation. It remains to be determined whether in these bivalent cells the OCT4:SOX2 accessible sites (and the genes they control) act as a break to differentiation. In line with this hypothesis, OCT4 has been suggested to prevent differentiation of pluripotent cells to trophectoderm by repressing CDX2 activity ^49^. Similarly, a dominant negative form of SOX2 has been shown to induce trophectoderm differentiation of mESCs, suggesting that SOX2 prevents exit from pluripotency ^50^. These mechanisms could also be relevant to prevent differentiation of progenitors in gastruloids.

The gastruloids derived from ΔCDX mESCs lacked NMPs, but still had a limited number of spinal cord cells. The presence of these spinal cord cells in absence of NMPs suggests that cell differentiation is not following rigid trajectories but may undertake alternative paths when the canonical ones are defective. This alternative path and therefore the origin of the spinal cord cells in ΔCDX gastruloids is still unclear. One possibility is that they are directly derived from the progenitor pool thus bypassing the NMP stage. Alternatively, as CDX genes are required for NMP state maintenance ^31^, the spinal cord cells in ΔCDX gastruloids may derive from NMPs that were present at an earlier stage during development but have disappeared at the final time point which prevents them to substantially contribute to the spinal cord population (Figure 4M).

Based on our pseudotime analyses, we did not find NMPs at the base of the spinal cord/mesoderm fork, as previously suggested ^4^. Rather we imputed that NMPs give origin to spinal cord cells, while the mesoderm lineages arise directly from the progenitor pool. This is line with a recent single cell transcriptomic analysis that proposes the presence of both NMP-dependent and NMP-independent developmental trajectories in gastruloids ^48^. However, we do not detect any clear NMP-dependent trajectory towards somitic mesoderm in the sciATACseq. These *in vitro* findings are in stark contrast with a large body of evidence suggesting that *in vivo* NMPs are bipotent cells capable of forming both spinal cord and paraxial mesoderm ^25,31^. Nonetheless if, as our data suggests, the lineage-restricted NMPs found in gastruloids are equivalent to NMPs in the embryo, it would imply that NMPs *in vivo* are not a homogeneous population but are rather represented by specialized subpopulations that can either differentiate to spinal cord or mesoderm cells. Consistent with our predictions, ΔCDX gastruloids were still able to form paraxial mesoderm in abundance. One possibility is that a subset of the gastruloid progenitor pool, which shows enrichment for SOX2 and TBXT motifs, represents an NMP-like population that does not match with E8.25 cells but could represent earlier embryonic NMPs. Our finding that ΔCDX gastruloids do not axially elongate recapitulates the phenotype of *Cdx* null mice, which fail to generate posterior tissues ^33^. NMPs in gastruloids are not only necessary to generate spinal cord cells but are essential for posterior end polarization, likely by the formation of a signaling center that secretes both Wnt and Fgf ligands ^33,51^.

By analyzing a mutant of the main driver of the paraxial mesoderm trajectory, we demonstrated that the intrinsic role of MSGN1 is to promote the formation of paraxial mesoderm in gastruloids. (Figure 4M) This finding is in line with *in vivo* studies showing that in mice *Msgn1* deletion leads to paraxial mesoderm loss ^45,46^. Our observation of TBXT+ cells in ΔMSGN1 gastruloids raises the possibility that these cells may be blocked in their differentiation path to paraxial mesoderm. Consistent with this, MSGN1 has been suggested to repress WNT3A signaling, which in turn promotes TBXT expression ^46,52^. Indeed, MSGN1 has been shown to repress TBXT expression. This repression could be required to progress into paraxial mesoderm, since MSGN1 null embryos present increased TBXT expression in immature paraxial mesoderm progenitors ^45^. Our pseudotime analysis suggests that, similar to paraxial mesoderm, the pharyngeal mesoderm originates from somitic mesoderm. Strikingly, we see that in ΔMSGN1 gastruloids there is an increase in pharyngeal mesoderm, indicating that MSGN1 suppresses pharyngeal mesoderm formation at the advantage of paraxial mesoderm differentiation (Figure 4M). Moreover, when we aimed to generate gastruloids from ΔMSGN1 ESCs, the resulting structures completely lacked spinal cord cells as well, which is in contrast with our prediction of MSGN1 driving only the mesoderm branch (Figure 4M). The use of chimeric WT/ΔMSGN1 gastruloids enabled us to disentangle the cell-intrinsic versus the cross-lineage roles of MSGN1. The fact that ΔMSGN1 cells, in the presence of WT-derived paraxial mesoderm, are able to efficiently differentiate into spinal cord cells shows that the communication between the paraxial mesoderm lineage and the spinal cord lineage is crucial for the formation of the latter (Figure 5H). Importantly, our observation that paraxial mesoderm induces spinal cord development is partially supported by observations in MSGN1 null mice which lack paraxial mesoderm and display a kinked neural tube accompanied with a reduction of SOX2 expression in the tail bud region of the embryo ^46^. However, the fact that MSGN1 null mice present only mild defects in neural tube formation might indicate that in the embryo the anterior tissues compensate for the absence of the paraxial mesoderm. Thus, CRN properties that are not evident in the complexity of the embryo can be brought to bear by the limited cell type composition of the gastruloid model.

CRNs cannot be studied in simple monocultures, since they require, by definition for multiple cell types to interact. However, within multicellular structures such as developing embryos and gastruloids perturbations to a GRN can have unexpected consequences as we have seen *in vitro* following the loss of MSGN1. Therefore, chimeric gastruloids, and in the future chimeric synthetic embryos ^53–55^, can serve as versatile tools to dissect and validate intrinsic versus extrinsic properties of regulators within a GRN that promotes tissue development. Our study serves as an initial framework for discovering how GRNs and CRN interact during development and lays the groundwork for future advances in understanding tissue interactions.

### STAR Methods

#### Cell lines

##### mESC culture

In this study we utilized E14 (IB10 subline) and HM1 mESCs, both derived from 129 mice ^56^. HM1, HM1-ΔCDX, HM1-ΔMSGN1 were cultured in 2i-LIF medium with 50% DMEM/F12 (Gibco) and 50% Neurobasal media (Gibco) supplemented with 0.5x N2 (Gibco), 0.5x B27 + retinoic acid (Gibco), 0.05% BSA (Gibco), 3μM CHIR99021 (BioConnect), 1μM PD03259010 (BioConnect), 2mΜ Glutamine (Gibco), 1.5×10-4 M 1-thioglycerol (Sigma-Aldrich), 100U/ml LIF (Cell guidance systems) and 50U/ml penicillin-streptomycin (Gibco) on gelatin-coated 10cm petri dishes in a humidified incubator (5% CO2, 37ºC). E14 cells have been derived from the Netherlands Cancer Institute, while HM1, HM1-ΔCDX and HM1-ΔMSGN1 cells were obtained from the lab of James Briscoe (Francis Crick Institute) ^31^.

##### Fluorescent protein labeling of mESCs

For the chimeric gastruloids experiments, E14, HM1-ΔCDX and HM1-ΔMSGN1 were genetically labeled with either EGFP or mCherry as indicated. The labeling was carried using a PiggyBac vector (kindly gifted by Bas van Steensel) ^57^ in which the coding sequence of EGFP or mCherry was cloned together with an EF1α promoter sequence. After transfection the positive cells were sorted by flow cytometry. The insert was designed *in silico* and synthesized by Twist Bioscience using Gibson assembly ^58^.

##### Gastruloid culture

Gastruloids were cultured as previously described ^3^. Briefly, gastruloids were aggregated in differentiation medium (N2B27), which includes DMEM/F12 and Neurobasal medium supplemented with 0.5X N2, 0.5X B27 + Retinoic acid, 2mΜ Glutamine, β-mercaptoethanol and 50U/ml penicillin-streptomycin in U-bottom 96-well plates (Thermo Scientific) in a humidified incubator (5% CO2, 37ºC). Cells were dissociated in N2B27 medium using a serological pipette combined with a 200µL pipette tip. Next, 3.75×10^4^ cells were transferred in N2B27 medium with final volume 5ml and 40µL of this suspension (∼300 cells) was transferred to each well. At 48 hours, 150µL of N2B27 medium with 3μM CHIR99021 were added using and Hamilton Star R&D liquid handling platform. From the day after, the medium was replaced with fresh N2B27 medium daily. For the chimeric gastruloid experiments, half of the total cell number was used from each line. Pictures were taken using an inverted wide field microscope (Zeiss Axio Observer Z1 Live) at 72, 96 and 120 hours. Gastruloids were fixed with 4% formaldehyde (Sigma-Aldrich) at approximately 120 hours, washed with PBS once and stored in PBS at 4ºC.

##### Cloning of pETNKI-6xhisSUMO3-Tn5

The gene encoding Tn5 was cloned into the pETNKI-6xhis-SUMO3-LIC vector, containing an N-terminal 6xHis-SUMO3-tag, using Ligation Independent Cloning (LIC) ^59^. The Tn5 gene was PCR-amplified using forward primer Tn5Fw 5’-ccagcagcagacgggaggtATGATTACCAGTGCACTGCATCGTGCGGCG -3’ and reverse primer Tn5Rv 5’ - ggcggcggagcccgttaGATTTTAATGCCCTGCGCCATCAGGTCTTTCGC -3’. pTXB1-Tn5 plasmid was a gift from Rickard Sandberg (Addgene plasmid #60240; http://n2t.net/addgene:60240; RRID: Addgene_60240) ^60^ and has been used as template DNA in the PCR reaction.

##### Expression of 6xhis-SUMO3-Tn5 protein

The recombinant Tn5 protein was expressed in Rosetta2 (DE3) cells in 3 liters of LB medium, supplemented with 30µg/ml kanamycin and 40 µg/ml chloramphenicol. Cells were grown at 37°C to OD_600_ = 0.6-0.8. the cell suspension was cooled to 18°C before 0.4 mM IPTG was added and protein was expressed overnight. Cells were harvested by centrifugation (3,000g, 15min, 20°C) and pellet was stored at -20°C until further use.

##### Tn5 protein purification

The Tn5 protein was essentially purified as described ^61^. Frozen cell pellet was resuspended in lysis buffer (20 mM Hepes pH 7.5, 800mM NaCl, 1 mM EDTA, 5% glycerol, 1 mM TCEP) containing 0.2% Triton-X100. After thawing, cells were lysed by sonication and the lysate was clarified by centrifugation (50,000g, 30min, 4°C). Polyethylenimine (0.1% w/v) was added dropwise to the supernatant, incubated for 30 min and the precipitate was removed by centrifugation (50,000g, 30min, 4°C). The soluble fraction was used for nickel affinity purification (1 ml Nickel Sepharose Excel, Cytiva). Beads were washed with lysis buffer, containing 20 mM imidazole, and protein was eluted by 200 mM imidazole in lysis buffer. Fractions were analyzed by SDS-PAGE. 6xHis-SUMO3-Tn5 protein was incubated with 6xHis-Senp2 protease to cleave off the 6xhis-SUMO3-tag, followed by overnight dialysis at 4°C against 20 mM HEPES pH 7.5, 800 mM NaCl, 1 mM EDTA, 5% glycerol, 1 mM TCEP. 6xHis-Senp2, 6xHisSUMO3 and residual intact 6xHis-SUMO-Tn5 was removed during reverse affinity purification using 1 ml Nickel Sepharose (Cytiva). The flowthrough was concentrated to 1 ml and the protein was further purified by size exclusion chromatography on a SEC 650 10/300 column (Bio-Rad), equilibrated with 20 mM HEPES pH 7.5, 800 mM NaCl, 1 mM EDTA, 5% glycerol, 1 mM TCEP. Fractions containing the monomeric Tn5 were pooled, concentrated and flash-frozen in liquid nitrogen before storage at -80°C.

##### Transposome formation

For both bulk and single cell ATAC experiments, transposons were annealed by mixing 10µL of 10X TE buffer with 45 µL of 100µM pMents oligonucleotides with 45 µL of 100µM of the corresponding i5 and i7 oligonucleotides (see oligonucleotides list for details). Next the adapter solution was incubated at 95ºC for 10min and cooled down to 4ºC at 0.1ºC/sec. For trasposome formation, the annealed adapters were diluted in equal volume of H_2_O. This adapter solution was mixed with 0.2mg/mL Tn5 in a 1:20 ratio and incubated for 1hr at 37ºC. The transposomes were either directly used for tagmentation or stored at - 20ºC for no longer than 2 weeks.

##### sciATACseq

sciATACseq was performed as previously described ^62^ with a few important adjustments. In particular, we implemented steps to reduce mitochondrial reads based on the Omni-ATAC protocol ^63^. In brief, 2-5×10^5^ cells were lysed for 3min in 500µL of RSB buffer (Tris-HCl pH 7.5 10mM, NaCl 10mM, MgCl_2_ 3mM, NP40 0.1%, Tween-20 0.1%, digitonin 0.01%) in presence of cOmplete protease inhibitor cocktail (Roche). Next, cells were resuspended 3 times and washed in 10 mL of RSBT buffer (Tris-HCl pH 7.5 10mM, NaCl 10mM, MgCl_2_ 3mM, Tween-20 0.1% and protease inhibitor cocktail) by inverting the tube 3 times and spinning at 500g for 10min at 4ºC. Next, the buffer was removed and 2500 nuclei were resuspended in 7.6µL of cold PBS and pipetted in a well of a 96-well plate with 1µL of i7 and i5 transposons and 10.4μL of transposition mix containing 10μL of 2xTD buffer (20 mM Tris, 10 mM MgCl2, 20% dimethylformamide, pH adjusted to 7.6), 0.2µL digitonin 1% and 0.2µL Tween 10%. The tagmentation was carried at 55ºC for 30min, nuclei were pooled and stained with 3μM 4′,6-diamidino-2-phenylindole (DAPI). The reaction was stopped by incubation at 37ºC for 15 min with 20μL 40 mM EDTA supplemented with spermidine 1mM. Next, 25 nuclei were sorted into either one or four 96 well plates (see run details) containing 11µL of EB buffer (Qiagen) and 1μL of Proteinase K 10 mg/ml (Roche). Decrosslinking was carried overnight at 65ºC and PCR was performed by adding 25µL of KAPA HiFi HotStart ReadyMix (Roche), 5μL of P5 and P7 primers (see oligonucleotides list)., 1μL of BSA 10mg/mL and 2 μL H_2_O. The thermocycler was set up as follows: 72ºC for 3min, 98ºC for 30s, 21 cycles of 98ºC for 10s, 63ºC for 30s,72ºC for 1min and finally 72ºC for 5min. Next, all PCR reactions were pooled and 1mL of the solution was purified using DNA clean & concentrator 5 (Zymo Research) following manufacturer instructions. Next, the library was size selected by 1x and 0.55x selection steps using AMPure XP beads (Beckman Coulter), followed by purification with QIAquick PCR purification kit (Qiagen). Finally, libraries were sequenced with a NextSeq 550 system (Illumina) using a customized protocol ^64^ (genomic DNA read 1 (cycles 1–51), index 1 (transposon i7, cycles 52–59, followed by 27 dark cycles, and PCR i7, cycles 60–67), index 2 (PCR i5, cycles 68–75, followed by 21 dark cycles, and transposon i5, cycles 76–83), and genomic DNA read 2 (cycles 84–134).

##### Bulk ATACseq

In brief, 50,000 mESCs were collected in cold PBS and lysed with a 2x lysis buffer (Tris-HCl pH7.5 1M, NaCl 5M, MgCl2 1M, 10% IGEPAL). Next, cells were pelleted and incubated with 2xTD buffer and 2 uL transposon mix. Next, PCR amplification was carried twice by KAPA HiFi HotStart ReadyMix using P5 and P7 indexed primers (see oligonucleotide list). Fragments between 200 and 700 bp were purified using AMPure XP beads. Quality control was performed by Bioanalyzer High Sensitivity DNA analysis (Agilent).

##### Analysis

###### Data preparation

Fastq files from single cell combinatorial indexing ATACseq were edited such that the combination of i5, i7, P5 and P7 barcodes (further referred to as ‘barcode’) from fastq comments were prepended with the ‘BC:Z:’ tag to allow transfer of barcodes to alignment files. Edited fastq files were mapped to the mm10 reference genome with the bwa program version 0.7.17-r118 using the command ‘bwa mem -MC’ ^65^. Mapped reads were filtered with samtools version 1.10 using ‘samtools view -h -b -q 10’ ^66^. Paired-end alignments were further processed to apply the +4, -5 Tn5 shift to proper pairs, deduplicated and matched to valid experimental barcodes, and resulting fragments were written to tabix files, using a custom R script. Fragments were considered duplicates when they matched chromosome, start and end positions, strand of the first mate and barcode.

###### Quality control

Barcodes were considered to represent cells when at least 2.000 unique nuclear fragments were represented by that barcode and the TSS score for that barcode exceeded 4. TSS scores were calculated by taking 100 basepairs centred at the transcription start site of UCSC’s known genes ^67^, along with 1000 basepair upsteam and downstream flanks. The number of overlapping fragments were determined for these TSSs and flanks, and divided by the 100 and 1000 basepair widths of the regions. The TSS score is then the ratio between the number for TSSs and the number for flanks. The fragments of included cells were then used to call peaks using MACS2 callpeak version 2.2.7.1 ^68^ with the arguments ‘-g mm -f BED –nomodel –extsize 200 –shift -100 –keep-duplicates=all’. A count matrix was constructed by quantifying the number of overlapping fragments in each cell with the peaks. Doublets were called with the scDblFinder ^69^ R package with the arguments ‘aggregateFeatures = TRUE, nfeatures = 25, processing = “normFeatures”’, which is their recommended setting for single cell ATACseq. We further excluded cells that had less than 2.000 fragments in peaks. In addition, we inspected banding scores ^19^, duplication rates, fraction of reads mapping to mitochondria, cell-wise GC bias, fragment length distributions and fraction of reads in peaks, but none of these were used as criteria for including or excluding cells.

###### Dimensionality reduction

From the binarized count matrix, peaks were excluded where fragments were observed in less than 2% of cells. Moreover, we excluded peaks that were within 1.5kb of a TSS ^70^ (exclude promoter bit). Term frequency – inverse document frequency (TF-IDF) with a log transformation on the TF ^19^, term was used as a weighting scheme for the binarized count matrix. The irlba package ^71^ was used to perform partial singular value decomposition of the scaled and centered TF-IDF matrix for the first 100 left and right singular vectors. The diagonal matrix was multiplied by the left singular vectors to obtain principal components. The mutual nearest neighbour method, per the ‘reducedMNN’ function in the batchelor R package ^26^, was used on the principal components to integrate cells from different batches. The 2^nd^-50^th^ corrected principal components were used to compute UMAPs using the uwot R package ^72^ with the arguments ‘metric = “cosine”, init = “agspectral”’.

###### Clustering and marker detection

Clustering of cells was performed using the Leiden algorithm on a shared nearest neighbour (SNN) graph computed on the batch-integrated principal components. The SNN graph was computed using the ‘makeSNNGraph’ function from the bluster R/Bioconductor package ^73^ with the argument ‘k = 15’. The ‘cluster_leiden’ function from the igraph ^74^ R package was used for clustering, with the ‘resolution_parameter = 0.1’ argument. Marker genes were detected using logistic regression to predict cells that had a fragment overlapping a peak from the total reads in peaks for a cell, cell-wise GC bias, batch membership and cluster membership. 20% of cells in each cluster were randomly combined to form a reference level for the cluster membership variable. In this regression, when the cluster membership had a significant log-odds ratio as determined by a Wald test after adjusting for false discovery rate, the peak was considered a marker peak for the tested cluster.

###### Motif analysis

Position frequency matrices of motifs were obtained with the ^75^ R/Bioconductor package from the vertebrates taxonomical group and core collection. For analysing motifs, we looked at the DNA content of peaks where cluster membership was a significant factor according to a likelihood ratio test, comparing the logistic regression model described above against a reduced model where cluster membership was omitted from the predictor variables. Peaks were then scanned for motif matches using the ‘matchMotifs’ function from the motifmatchr ^76,77^ R/Bioconductor package. Motif Z-scores stabilised for GC bias were then computed using the chromVAR R/Bioconductor ^78^ package.

###### Pseudotime analysis

Diffusion pseudotime was calculated with the destiny R/Bioconductor package ^26^ using the ‘DiffusionMap’ function with the ‘distance = “cosine”’ argument, using the 2^nd^ to 50^th^ corrected principal components as data The pseudotime was extracted using the ‘DPT’ function from the destiny package. The cells in the endothelium and endoderm clusters were omitted from this computation due to their apparent discontinuity from other clusters. From the diffusion maps, it was inferred that differentiation largely occurred along two branches originating from the same progenitor cluster. Based on the diffusion maps, a decision was made to split cell into a spinal cord arm and mesoderm arm according to Figure S3A.

###### Comparison with embryo

Peaks were called as described above on combined fragments from both the embryo data and our gastruloids data. Cell annotations were taken from the supplementary data of ^14^. Fragments were counted in these peaks to generate a count matrix, and from this step the data was processed in the same way as described above, with the following exceptions. We required peaks to have fragments in 1.5% of cells instead of 2% in the dimensionality reduction step. During batch integration, the embryo data was treated as another batch. For learning labels from the embryo data, we trained a support vector machine (SVM) ^79^ on batch-integrated principal components to classify embryo cells using the e1071 R package. 60% of embryo cells were used as training data. 20% were used to tune the ‘gamma’ parameter in the ‘svm’ function, which was eventually set at 1e-4. The remaining 20% of embryo cells were used to test accuracy. The SVM reached an accuracy of 97.3% on this 20% of hold-out cells. Naturally, this accuracy reflects the performance on cells from the same dataset, and we can’t know how well this performance translates to our gastruloids data across datasets.

##### Morphological analysis

The morphology of gastruloids and the spatial distribution of fluorescently labelled cells were measured with the Axios Observer Z1 microscope (Zeiss). For the morphological scoring, wide field gastruloid pictures were scored blindly, and used to divide the gastruloids into ovoid, elongated and multipolar based on their morphology. For the ΔCDX gastruloids experiments, the axis ratio was measured using Fiji ^80^. To quantify the spatial distribution of the fluorescence signal along the anterior-posterior axis of the gastruloids, Fiji was used to draw a line from the anterior to the posterior end in order to measure the mean intensity of EGFP or mCherry along the axis.

##### Immunofluorescent confocal staining and analysis

The staining of the gastruloids was performed as previously described ^81^. After 30min fixation, 4% formaldehyde was washed away with PBS and gastruloids were blocked with PBS-FT, which consists of 10% fetal bovine serum and 0.2% Triton-X100 (Sigma Aldrich) in PBS for 1 hour at room temperature. Next, gastruloids were stained with the following primary antibodies diluted in PBS-FT: 1:100 goat-anti-SOX2 (Biotechne), 1:500 rabbit-anti-FOXC1 (Abcam) or 1:250 goat-anti-BRA (Biotechne), 1:100 rabbit anti-CDX2 (Cell Signalling Technology). Gastruloids were incubated at 4ºC overnight. After the incubation, gastruloids were washed with PBS-FT three times. For the secondary antibodies staining, 1:250 donkey-anti-rabbit 405 (ThermoFisher), 1:250 Donkey-anti-goat 488 (ThermoFisher), 1:250 Donkey-anti-rabbit 594 (Jackson) or 1:500 donkey-anti-goat 647 (Abcam) were added to the gastruloids together with 3μM DAPI where indicated and incubated at 4ºC overnight. Finally, the secondary antibodies were washed three times and the gastruloids were mounted in the microscope slides using FluorSafe (Sigma Aldrich). Confocal imaging was performed using the SP5 confocal microscope (Leica). For the ΔCDX and ΔMSGN1 gastruloid experiments, one confocal plane per gastruloid was imaged, while for the experiments including the chimeric gastruloids 3 subsequent confocal planes were imaged and averaged in the analysis. To investigate the spatial distribution of the different cell types or staining along the gastruloid, Fiji was used to segment the positive areas and measure the intensity and area in all the channels.

##### Statistical analysis

For the image quantification analysis, analysis of variance (ANOVA) was performed.

## Acknowledgements

We would like to thank members of the de Wit laboratory for critically reading the manuscript. We thank Alfonso Martinez Arias for insightful comments on a draft version of this manuscript. We thank James Briscoe for kindly providing CDX and MSGN1 knock-out mouse ESC lines. We would like to thank the facilities of the NKI, particularly the Flow Cytometry facility for assistance in sciATACseq experiments, the Research High Performance Computing facility for supporting the computational analysis, the Genomics Core facility for sequencing, the High Throughput Screening facility for gastruloid culturing automation and the BioImaging Facility for supporting microscopy analysis. Work in the de Wit lab is supported by ERC Consolidator grant (865459, ‘FuncDis3D’) and a Vidi grant ‘016.16.316’ from the Dutch Research Council (NWO). L.B., T.vd.B., N.A.S. and E.d.W. are part of Oncode with is partially funded by the Dutch Cancer Society.

