## Supplemental Figures for "Identifying cross-lineage dependencies of cell-type specific regulators in gastruloids"

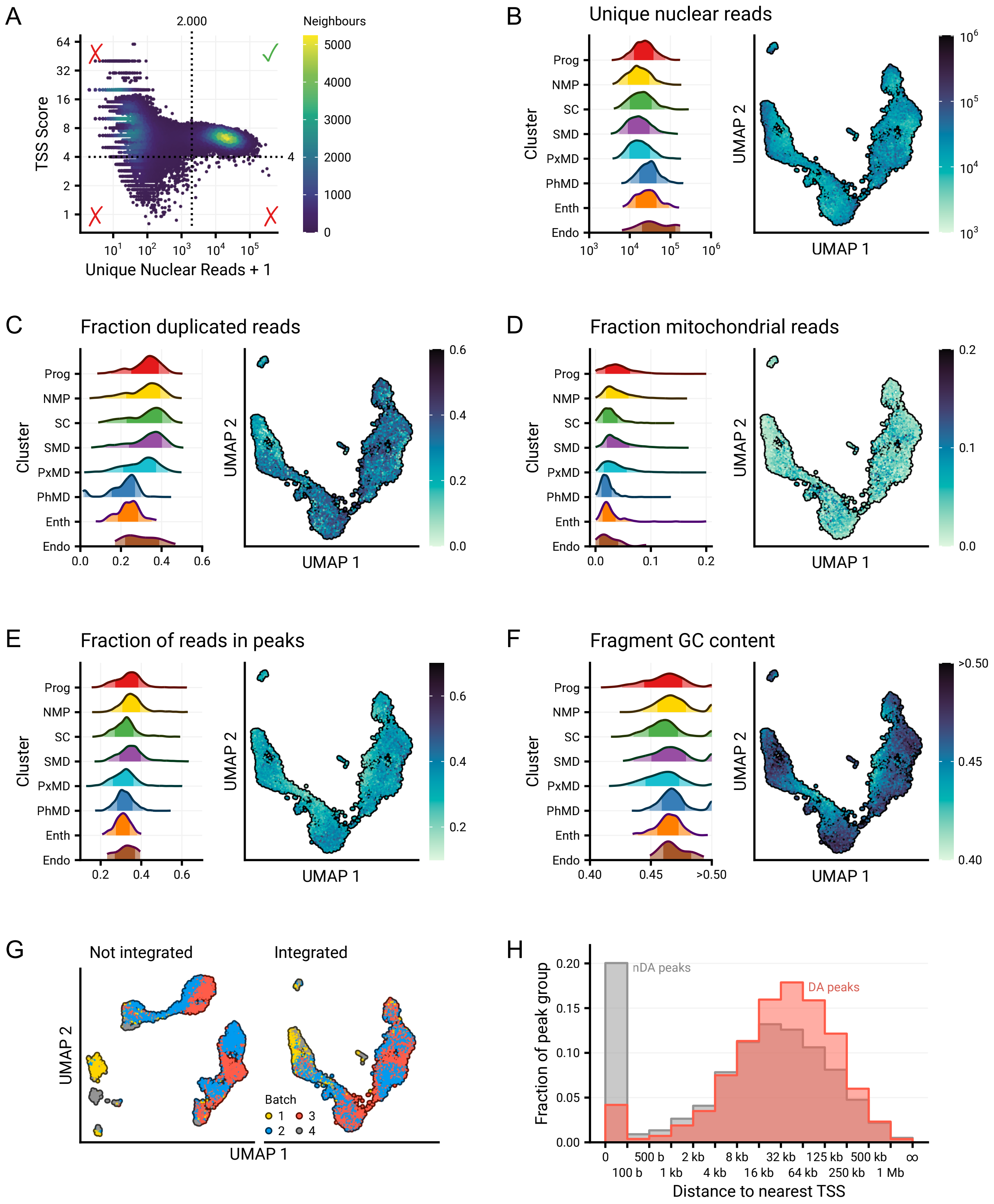
A
2.000
Neighbours
5000
4000
3000
2000
1000
0
TSS Score
Unique Nuclear Reads + 1
B
Unique nuclear reads
C
Fraction duplicated reads
D
Fraction mitochondrial reads
E
Fraction of reads in peaks
F
Fragment GC content
Prog
NMP
SC
SMD
PxMD
PhMD
Enth
Endo
Cluster
UMAP 1
UMAP 2
G
Not integrated
Integrated
Batch
H
nDA peaks
DA peaks
Fraction of peak group
Distance to nearest TSS
0
100 b
500 b
1 kb
2 kb
4 kb
8 kb
16 kb
32 kb
64 kb
125 kb
250 kb
500 kb
1 Mb

**Figure S1:**

Quality control metrics for sci-ATACseq. A) Inclusion criteria for considering barcodes as cells, wherein a cell needs at least 2.000 nuclear reads and have a transcription start site (TSS) score higher than 4 (see methods). Every point is a barcode, and local neighbour count is displayed as proxy for density. B-F) Quality control metrics displayed as ridge plots (left) and in relation to UMAP coordinates (right). Metrics are the number of unique nuclear fragments (B), the fraction of non-unique (duplicated) nuclear fragments out of total nuclear fragments (C), the fraction of unique mitochondrial reads out of total unique reads (D), the fraction of reads in peaks (E), the fraction of GC basepairs over all fragments in a cell (F). G) Comparison of UMAP dimensionality reduction before and after batch integration through the mutual nearest neighbours method. H) Histogram of distances between peaks and their nearest TSS compared between differentially accessible (DA, marker for any population) and not differentially accessible (nDA) peaks.

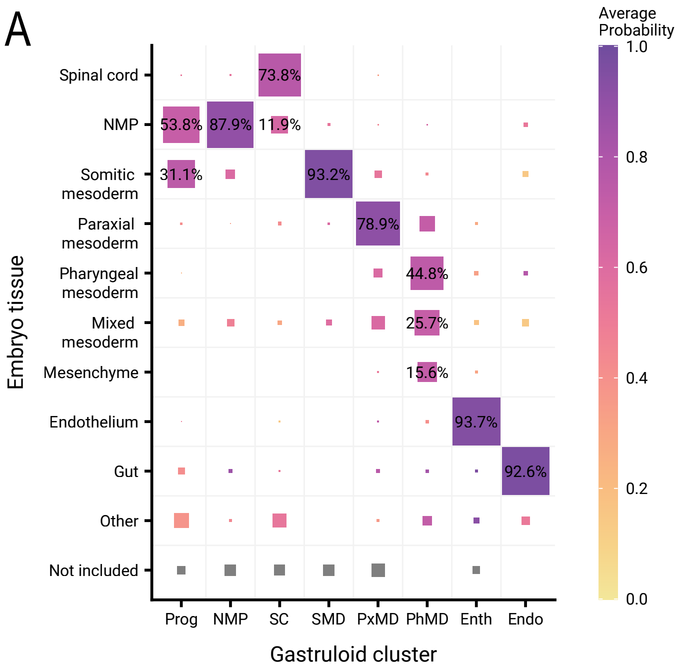
A
Average Probability
1.0
0.8
0.6
0.4
0.2
0.0
Embryo tissue
Spinal cord
NMP
Somitic mesoderm
Paraxial mesoderm
Pharyngeal mesoderm
Mixed mesoderm
Mesenchyme
Endothelium
Gut
Other
Not included
73.8%
53.8%
87.9%
11.9%
31.1%
93.2%
78.9%
44.8%
25.7%
15.6%
93.7%
92.6%
Prog
NMP
SC
SMD
PxMD
PhMD
Enth
Endo
Gastruloid cluster

**Figure S2:**

Support vector machine (SVM) trained by embryo cells predicts labels for gastruloid cells. A) Proportions of gastruloid cells belonging to clusters (x-axis) predicted as being embryo tissue (y-axis). Area of squares is proportional to the fraction of cells in the gastruloid cluster that have a specific prediction, whereas colour gives the average probability, according to the SVM, of this intersection. ‘Other’ indicates the sum of embryo tissues not displayed in this plot. ‘Not included’ indicates gastruloid cells that do not have more than 2.000 accessible peaks, as called on the combined gastruloid and embryo data.

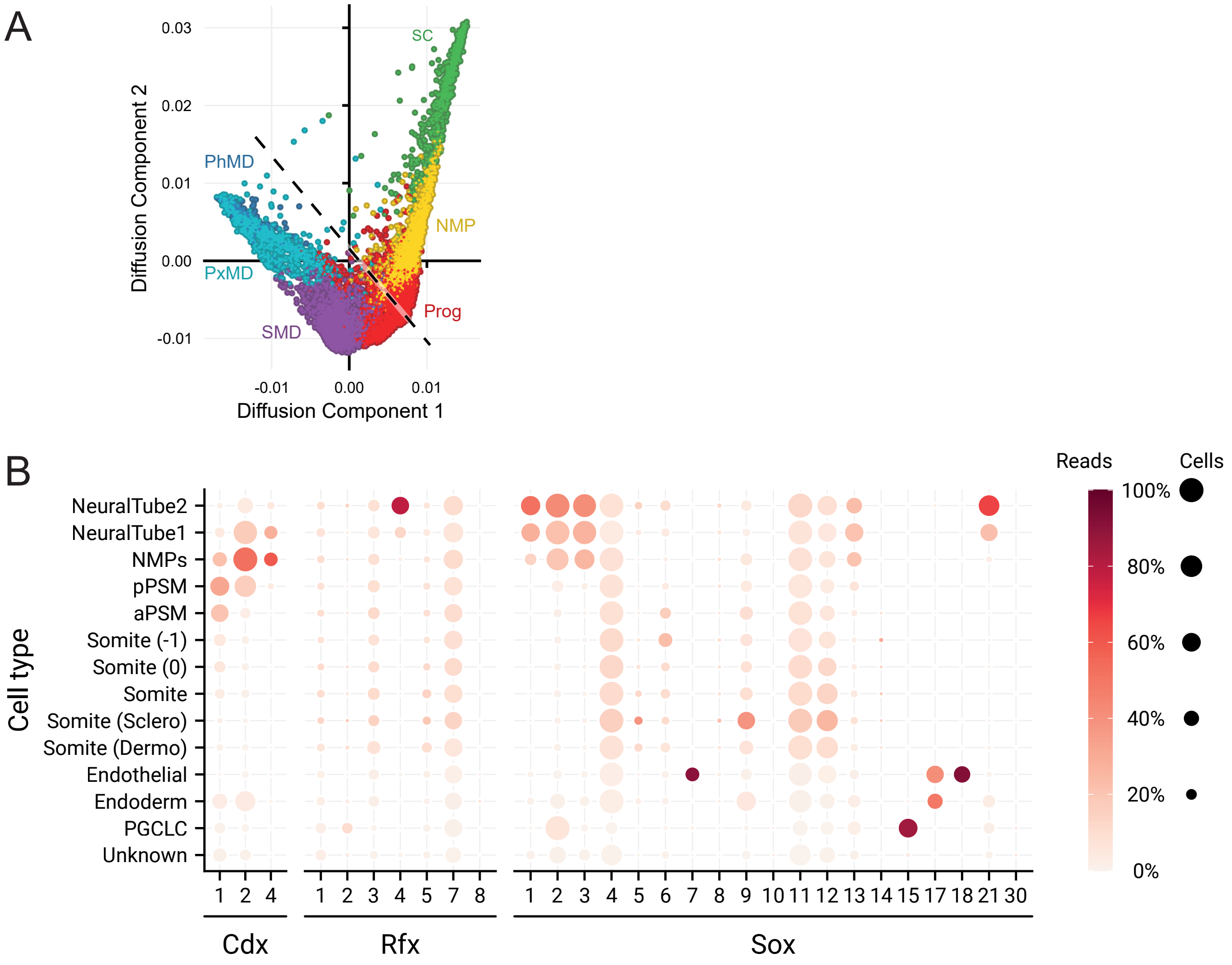
A
Diffusion Component 2
Diffusion Component 1
0.03
0.02
0.01
0.00
-0.01
-0.01
0.00
0.01
SC
PhMD
NMP
PxMD
Prog
SMD
B
Cell type
NeuralTube2
NeuralTube1
NMPs
pPSM
aPSM
Somite (-1)
Somite (0)
Somite
Somite (Sclero)
Somite (Dermo)
Endothelial
Endoderm
PGCLC
Unknown
Cdx
1 2 4
Rfx
1 2 3 4 5 7 8
Sox
1 2 3 4 5 6 7 8 9 10 11 12 13 14 15 17 18 21 30
Reads
Cells
100%
80%
60%
40%
20%
0%

**Figure S3:**

A) Scatterplot of diffusion components coloured by cluster label. Cells were divided into the spinal cord arm or mesdoderm arm based on the split indicated by the dashed line. B) Summary of gastruloid gene expression as measured by scRNA-seq data [4] for highly enriched motif families in the NMP-spinal cord branch. Bubble plot showing percentage of cells expressing a gene through bubble size, and percentage of reads of that gene allocated to a cell type through colour.

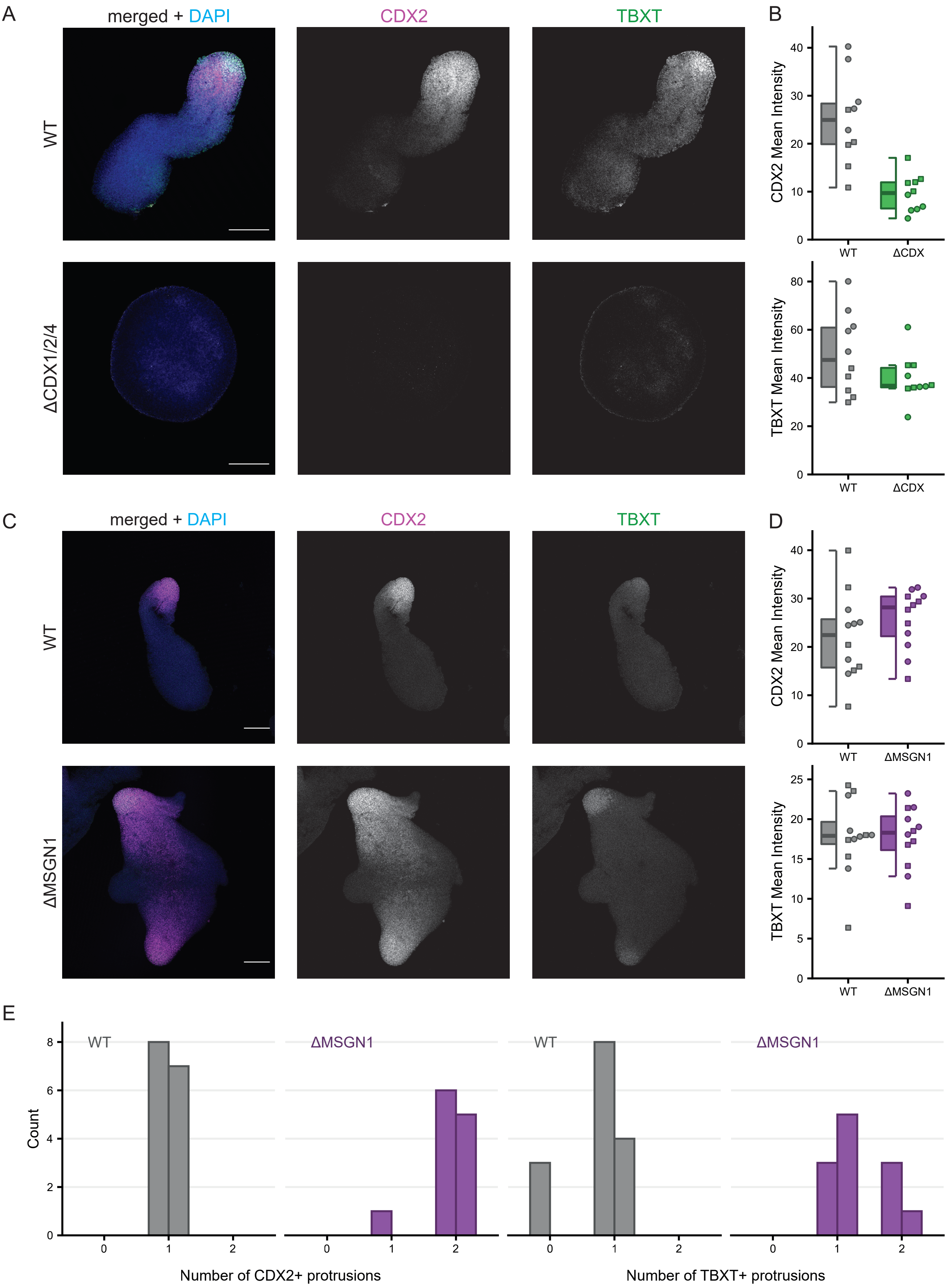
A
merged + DAPI
CDX2
TBXT
WT
ΔCDX1/2/4
B
CDX2 Mean Intensity
TBXT Mean Intensity
WT
ΔCDX
C
D
ΔMSGN1
E
Count
Number of CDX2+ protrusions
Number of TBXT+ protrusions

**Figure S4:**

CDX2 and TBXT expression in ΔCDX and ΔMSGN1 gastruloids. A) Confocal immunofluorescence images of WT and ΔCDX gastruloids at 120 hours, stained for CDX2 (magenta) and TBXT (green). DAPI in blue. Scale bar: 200 µm. B) Quantification of CDX2 and TBXT expressionin WT and ΔCDX gastruloids (n = 2 biological replicates; rep. 1 = circles, WT: 5 gastruloids, ΔCDX: 5 gastruloids; rep. 2 = squares, WT: 5 gastruloids, ΔCDX: 5 gastruloids). C) Like A, but for WT and ΔMSGN1 gastruloids (n = 2 biological replicates; rep. 1 = circles, WT: 6 gastruloids, ΔMSGN1: 6 gastruloids; rep. 2 = squares, WT: 6 gastruloids, ΔMSGN1: 6 gastruloids). D) Quantification of CDX2 and TBXT expression in WT and ΔMSGN1 gastruloids. E) Quantification of CDX2+ and TBXT+ protrusion counted in WT and ΔMSGN1 gastruloids.

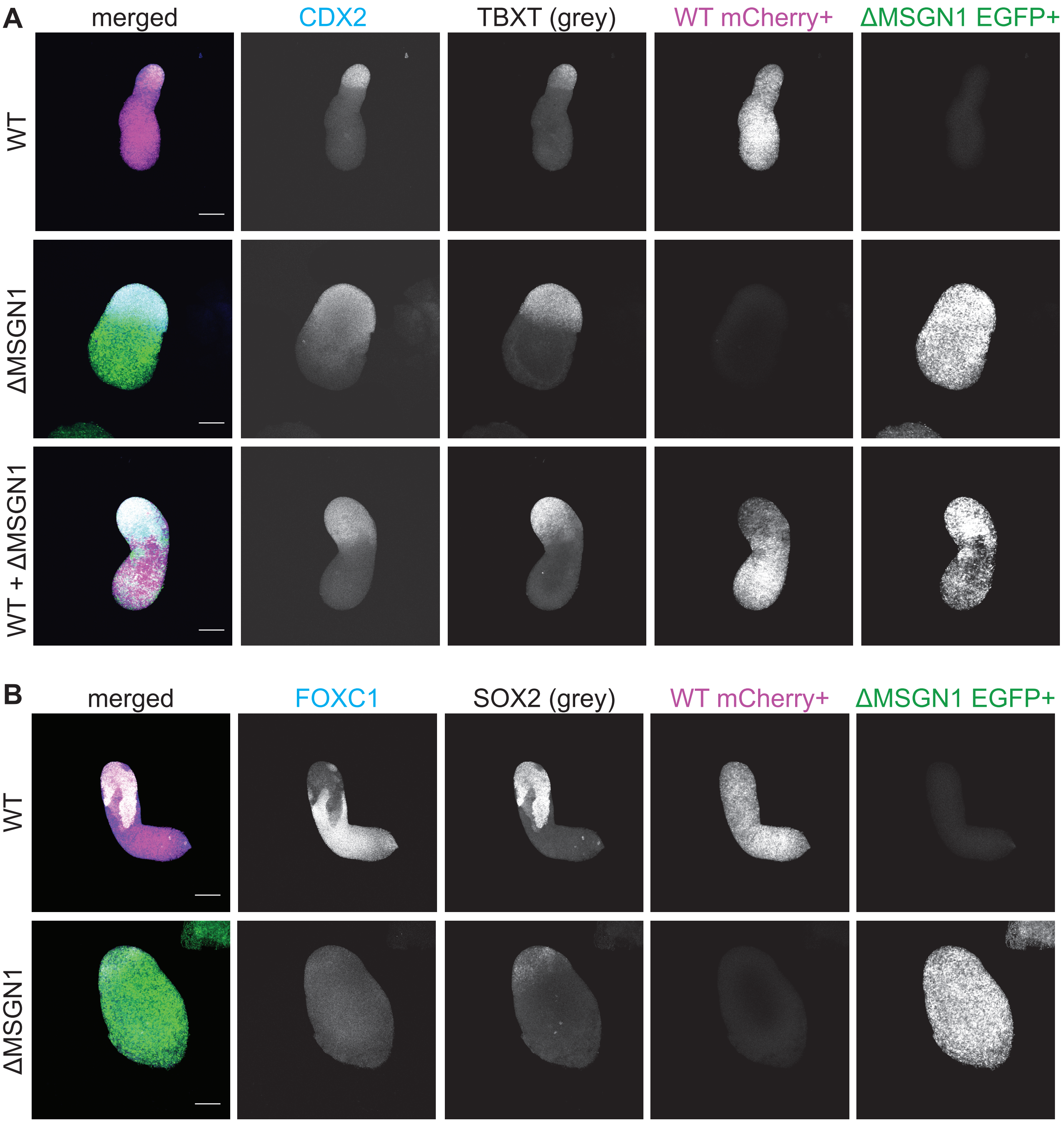
A
merged
CDX2
TBXT (grey)
WT mCherry+
ΔMSGN1 EGFP+
WT
ΔMSGN1
WT + ΔMSGN1
B
merged
FOXC1
SOX2 (grey)
WT mCherry+
ΔMSGN1 EGFP+
WT
ΔMSGN1

**Figure S5:**

Lineage marker expression in chimeric WT + ΔMSGN1 gastruloids. A) Confocal immunofluorescence images of WT only, ΔMSGN1 only and chimeric WT + ΔMSGN1 gastruloids at 120h, stained for CDX2 (blue) and TBXT (grey), mCherry (magenta) and EGFP (green). Scale bar: 200 µm. B) Confocal immunofluorescence images of WT only and ΔMSGN1 only gastruloids at 120h, stained for the FOXC1 (blue), SOX2 (grey), EGFP (green) and mCherry (magenta).
